# Grb7 is dispensable for Erbb2-driven mouse mammary tumorigenesis

**DOI:** 10.1101/2025.06.15.659799

**Authors:** Kristopher A. Lofgren, Peter Feiszt, Paraic A. Kenny

## Abstract

Growth factor receptor-bound 7 (Grb7) is a multidomain adaptor protein implicated in signal transduction from multiple receptor tyrosine kinases, including ERBB2. *ERBB2* amplification is a common event in breast cancer, and co-amplification of the neighboring *GRB7* gene typically occurs, which has been presumed to lead to synergistic pro-tumorigenic signaling between both encoded proteins. Accordingly, GRB7 has been proposed as a candidate therapeutic target in breast cancer and other malignancies.

Genetic deletion of *Grb7* results in relatively phenotypically normal, viable, fertile mice. The sole defect observed in these animals was a mammary dysfunction resulting in a failure to efficiently nurse pups to weaning. Here we sought to directly evaluate the extent to which Grb7 expression may be required for Erbb2-driven mammary tumorigenesis by crossing these *Grb7* knockout mice with MMTV-*Neu* transgenic mice. Both *Grb7* deficient and proficient MMTV-*Neu* cohorts developed tumors at very similar rates, demonstrating that Grb7 is dispensable for Erbb2-driven tumorigenesis in this model.

## INTRODUCTION

Growth factor receptor-bound 7 (Grb7) is a multidomain adaptor protein implicated in the regulation of cell proliferation, survival, migration, and invasion through its interacting proteins, which include EGFR/ErbB2/ErbB3, Shc, Ret, PDGFR, FGFRs, EphB1, c-Kit, FAK, Tek/Tie2, insulin receptor, caveolin, and calmodulin (reviewed in Chu et al., 2019; Han et al., 2001). In breast cancer, *GRB7* was shown to be co-amplified with *ERBB2* (Stein et al., 1994). GRB7 has been associated with poor outcomes in breast cancer case series (Nadler et al., 2010; Ramsey et al., 2011) and is one of the 21 genes used to compute the Oncotype Dx recurrence score (Sparano et al., 2015). GRB7 inhibitors have been developed (Ambaye et al., 2011; Tanaka et al., 2006; Watson et al., 2017), with activity in pre-clinical breast cancer models (Giricz et al., 2012; Pero et al., 2007). GRB7 expression is also altered in several other cancer types (Chu et al., 2019). In aggregate, data suggest that GRB7 may contribute to tumorigenesis via its role in receptor tyrosine kinase signaling, with a potential role in transducing ERBB2 signaling in breast cancer being a leading contender, however this has not been experimentally evaluated in a genetic model.

The cytoplasmic phospho-tyrosines of active ERBB2 recruit several different adaptor proteins which orchestrate multiple downstream signaling cascades. GRB7 was initially demonstrated to strongly and directly bind to both ERBB2 and Shc (Stein et al., 1994). GRB7 directly binds the ERBB2 p-YB site, corresponding to Y1139 in Uniprot accession P04226, (Janes et al., 1997), a site that it shares with GRB2 (Dankort et al., 1997). Shc binds the pY-D/E site (Y1221/1222) and, in turn, can recruit GRB2 or GRB7 in a mutually exclusive manner (Stein et al., 1994). Accordingly, GRB2 and GRB7 may be recruited to ERBB2 both directly and indirectly, and may also be recruited to ERBB2 heterodimerization partners such as EGFR (Margolis et al., 1992). The conditional deletion of *Shc* completely ablated *in vivo* mammary tumorigenesis driven by an activated *Erbb2* (*Neu*) transgene (Ursini-Siegel et al., 2008), indicating a strong requirement for signaling via the pY-D/E site for mammary tumorigenesis *in vivo* and suggesting that, in the absence of Shc, recruitment of either Grb2 or Grb7 directly to the pY-B site of ERBB2 is not sufficient to efficiently promote tumorigenic signaling.

We recently generated *Grb7* knockout mice (Lofgren and Kenny, 2024). Somewhat surprisingly, these mice lack gross developmental abnormalities and were viable and fertile. The sole phenotype observed was the inability of knockout females to support pups to weaning age, implicating Grb7 in mammary function. Two independently generated *Grb7* knockout models also show modest, primarily metabolic, phenotypes (Moorwood et al., 2024; Vermehren-Schmaedick et al., 2024). Given the extensive literature associating ERBB2 and GRB7 in cancer, the present study evaluates the extent to which Grb7 expression is required for ERRB2-driven mammary tumorigenesis in the MMTV-Neu model (Guy et al., 1992).

## MATERIALS AND METHODS

### Mouse strains and cohort generation

All animal use and procedures were approved by the University of Wisconsin – La Crosse Institutional Animal Care and Use Committee. *Grb7* knockout mice on a C57BL/6 background were previously described (Lofgren and Kenny, 2024) and maintained in-house. MMTV-Neu mice (FVB/N-Tg(MMTVneu)202Mul/J) were obtained from Jackson Laboratories (Strain #002376)(Guy et al., 1992). Mice were housed in individually-ventilated cages on a 12-hour light/12-hour dark cycle. Food (Harlan 2018 rodent chow) and water provided *ad libitum*. Genotyping was performed with tail biopsy DNA using the following primers: *Grb7*: TGGGATTGGCATTTTGTCTG, TCGTGGTATCGTTATGCGCC, AAGCCAGTGTTCAGCCTCC; MMTV-Neu: TTTCCTGCAGCAGCCTACGC, CGGAACCCACATCAGGCC; MMTV-Neu zygosity check: GCAGAAGATGGCCTAGTCGG, TGTGTCCCTGAATGCAAGTTT, AACCATGTCTTTGAGTATAG.

*Grb7* KO (*Grb7*^*del*/del^) males were paired with MMTV-Neu homozygote (*Neu*^*Tg*/Tg^) females to generate a hybrid F1 generation of *Neu*^*Tg*/0^ *Grb7*^del/+^ mice with a 1:1 mixed background of FVB and C57BL/6. F1 dihybrid crosses were used to generate the F2 females for the study cohort of 25 *Neu*^*Tg*/0^*Grb7*^del/del^ and 44 *Neu*^Tg/0^*Grb7*^del/+^ mice. Nulliparous females were aged in group housing and were inspected weekly by palpation to detect tumor formation.

### Statistical Analysis

The primary study endpoint was time to palpable tumor detection. Data were analyzed using Kaplan-Meier curves and a log-rank p-value < 0.05 was considered statistically significant.

## RESULTS

Cohorts consisting of 44 MMTV-*Neu/Grb7*^+/-^ and 25 MMTV-*Neu/Grb7*^-/-^ nulliparous female mice were generated and followed for up to 142 weeks. Tumor time of onset was assessed by palpation and all palpated masses were confirmed as tumors by histological analysis after euthanasia. Median age of tumor palpation was 97.2 weeks for MMTV-*Neu*/ *Grb7*^+/-^ and 96.9 weeks for MMTV-*Neu*/ *Grb7*^-/-^ (Figure 1, P = 0.91). Accordingly, Grb7 was not required for Erbb2-driven tumorigenesis in this model. Few mice had multiple mammary tumors (5 *Neu*, avg multiplicity of 1.27; 3 *Neu*/*Grb7*KO avg multiplicity of1.35), which indicated *Grb7* status did not significantly alter tumor multiplicity. Full necropsies were performed on tumor-bearing mice to investigate the presence of metastatic disease. Four MMTV-*Neu*/*Grb7*^+/-^ and 1 MMTV-*Neu*/*Grb7*^-/-^ mouse exhibited lesions in the lungs or liver, which indicated no impact of Grb7 on tumor metastasis in this study.

**Figure 1.**
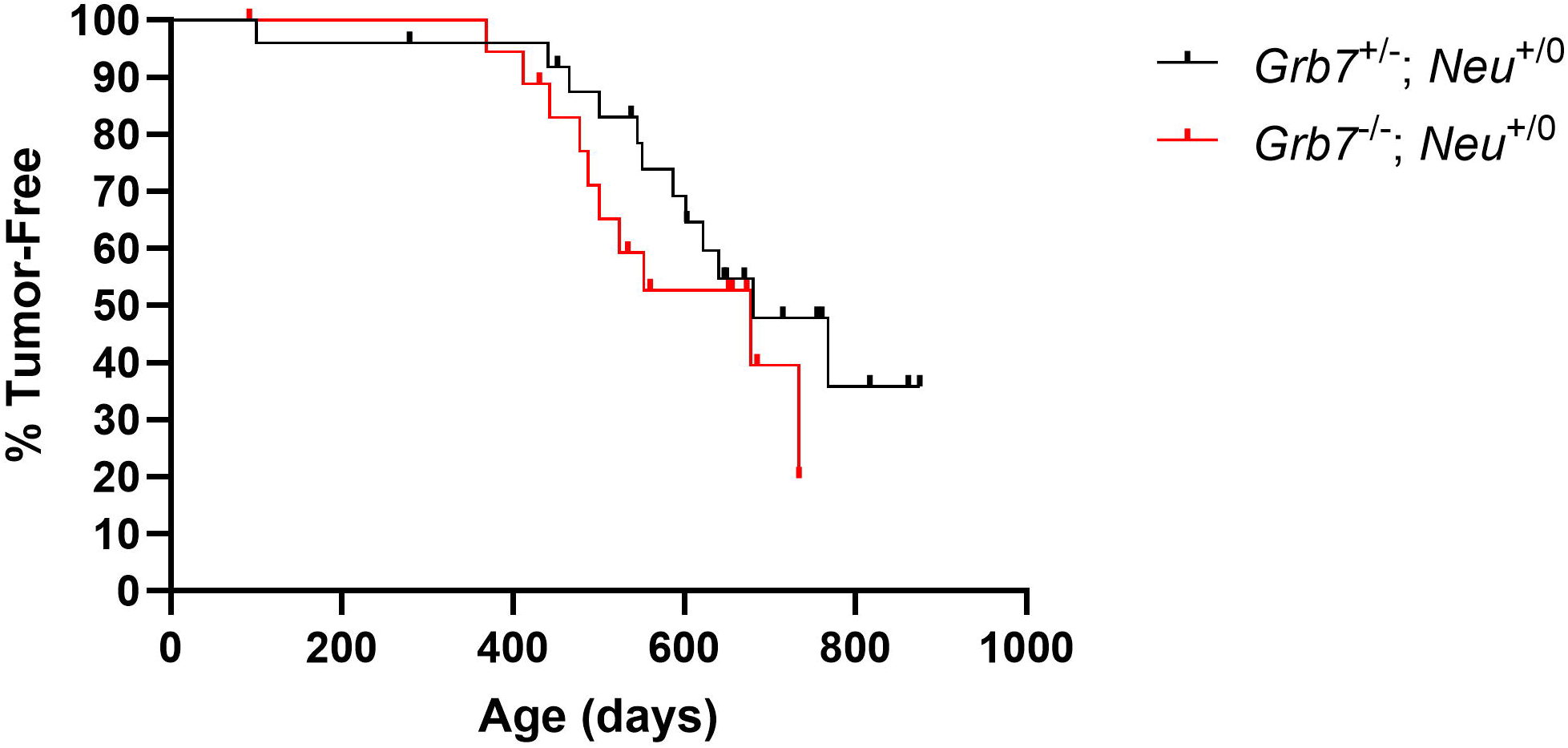
MMTV-*Neu* driven tumorigenesis is not affected by *Grb7* deletion. Mice were observed daily and palpated weekly. Time to tumor onset was not different between *Grb7* arms, with a mean time to detection of 681 days (97.2 weeks) for MMTV-*Neu*/*Grb7*^+/-^ mice and 678 days (96.9 weeks) for MMTV-*Neu*/*Grb7*^-/-^ mice; p = 0.91.

A prior study that added back individual tyrosines to a Neu transgene with all phospho-tyrosines ablated found that tumors driven by the pY-B and pY-D sites had different histologies (Dankort et al., 2001) suggesting that differential effector recruitment may favor different morphologies. In our cohort, hematoxylin and eosin staining showed a range of carcinoma histologies, including uniform solid, cribriform, sheet-like, or glandular-like organization. We evaluated whether *Grb7* deletion changed the relative proportions of these histologies, but detected no difference in tumor organization or composition between the two experimental arms.

## DISCUSSION

Our findings demonstrate that Grb7 is not required for ERBB2-driven mammary tumorigenesis, as evidenced by similar tumor onset times and multiplicity between *Grb7* knockout and heterozygous cohorts. This was unexpected given the extensive literature suggesting a role for Grb7 in propagating signals from ERBB2, as well as studies demonstrating that inhibitors of Grb7 could affect tumor phenotypes in both *in vitro* and *in vivo* models.

Demonstrating an attenuation in tumorigenesis in the *Grb7* knockout animals would have implied either or both of (1) an essential role for Grb7 at the pY-B site or (2) that recruitment of Grb7 to Shc at the pY-D site is essential for tumorigenic signaling. Because we found no difference in tumorigenesis rate, we can reject both propositions. Instead, it seems likely that potential redundancy with Grb2 might compensate for any deficit resulting from Grb7 loss.Future studies incorporating conditional deletion of *Grb2* would help clarify this point.

The lack of impact of *Grb7* deletion on ERBB2-driven tumorigenesis raises significant questions about the utility of Grb7 as a therapeutic target. A cell-penetrating peptide inhibitor of Grb7, G-718-NATE, was shown to inhibit proliferation of multiple breast cancer cell lines in vitro, and to synergize with trastuzumab in suppressing proliferation of the *ERBB2*-amplified SKBR3 cell line (Pero et al., 2007). Injection of this peptide also attenuated the growth of peritoneal metastases of injected Panc-1 pancreatic cancer cells in immunocompromised mice (Tanaka et al., 2006). While surface plasmon resonance data indicate a strong selectivity of this peptide for Grb7 over Grbs 2/10/14 (Gunzburg et al., 2012), it is entirely possible that the antiproliferative and antimigratory effects observed in these studies are mediated by blocking Grb7 interactions with proteins other than ERBB2. Indeed, our prior work with this peptide in triple-negative breast cancer cell lines (Giricz et al., 2012), demonstrated potent effects in an ERBB2 low-to-negative setting.

When comparing our data with studies demonstrating an effect of this inhibitor on *ERBB2*-amplified breast cancer cell lines (Pero et al., 2007), it is also important to consider that these were late-stage treatments of established ERBB2-driven tumors that developed in the presence of co-amplified *GRB7*. In our model, the ERBB2-driven tumors develop in the absence of Grb7 so it is possible that they adapt to this absence by favoring other signaling adaptors during tumor initiation. In this way, our data may not be directly comparable with these prior cell line studies and it is certainly possible that ERBB2-GRB7 interactions may be important when tumors develop with co-amplification of these neighboring genes. In that case, the excess production of Grb7 might lead to such a strong imbalance of Grb7 v Grb2 that Grb7 recruitment to ERBB2 is more strongly favored, potentially leading to a stronger dependence on Grb7 that was revealed by the cited inhibitor studies.

The very strong correlation between *GRB7* and *ERBB2* copy number-driven expression increases in breast cancer has been used to infer likely synergistic promotion of tumorigenesis, but this close genetic linkage complicates the determination of true causality. Accordingly, counterexamples showing *Erbb2*-driven tumorigenesis in the absence of *Grb7* alteration, though rare, may be quite informative. Tumors developing in the MMTV-*NeuNT* activated *Erbb2* (V664E) mouse model in which this transgene is knocked into the endogenous *Erbb2* locus typically gain Erbb2 amplification and have been used to map the co-amplified genes at that locus (Hodgson et al., 2005). As found in human breast cancer, *Erbb2* and *Grb7* are almost always co-amplified and co-overexpressed in the tumors formed in these animals. Interestingly, the authors described a single tumor (of 20 analyzed) that amplified *Erbb2* without *Grb7* co-amplification demonstrating, at least in the activated NeuNT model, that *Grb7* co-amplification is not essential for tumorigenesis. Notably, modest levels of Grb7 protein expression were detected in this tumor, so the study findings did not imply that Grb7 was entirely dispensable.

Genetic background and parity are both important influences on mammary tumor latency in transgenic mouse models (Henry et al., 2004; Jamerson et al., 2003; Lifsted et al., 1998) and are important to control for. Most studies on MMTV-*Neu* induced tumorigenesis have utilized the FVB background (reviewed in Simond and Muller, 2020). The *Grb7* knockout was developed in the C57BL/6 background, which is typically less susceptible to mammary tumorigenesis than FVB, making it especially important to control for potential contributions of strain-associated modifier alleles. Although we did not back-cross to a single background, the breeding scheme was controlled to result in female mice with a 50:50 contribution from each background so that the mixed background would be consistent across both *Grb7*^+/-^ and *Grb7*^-/-^ cohorts. To exclude any potential influence of parity on tumor onset time, all study cohort mice were non-parous.

In conclusion, our study demonstrates that Grb7 is not essential for tumorigenesis in the MMTV-Neu model, which may prompt a reevaluation of its potential as a therapeutic target for breast malignancies.

## ACKNOWLEDGEMENTS

This study was funded by the Gundersen Medical Foundation. PK holds the Dr. Jon & Betty Kabara Endowed Chair in Precision Oncology. KL was supported by the Norman L. Gillette, Jr. Cancer Research Fellowship.

## Notes

### Competing Interest Statement

The authors have declared no competing interest.

